# Dimensional clinical phenotyping using post-mortem brain donor medical records: Association with neuropathology

**DOI:** 10.1101/2023.05.04.539430

**Authors:** Jonathan S. Vogelgsang, Shu Dan, Anna P. Lally, Michael Chatigny, Sangeetha Vempati, Joshua Abston, Peter T. Durning, Derek H. Oakley, Thomas H. McCoy, Torsten Klengel, Sabina Berretta

## Abstract

**INTRODUCTION:** Transdiagnostic dimensional phenotypes are essential to investigate the relationship between continuous symptom dimensions and pathological changes. This is a fundamental challenge to postmortem work, as assessment of newly developed phenotypic concepts needs to rely on existing records.

**METHODS:** We adapted well-validated methodologies to compute NIMH research domain criteria (RDoC) scores using natural language processing (NLP) from electronic health records (EHRs) obtained from post-mortem brain donors and tested whether RDoC cognitive domain scores were associated with hallmark Alzheimer’s disease (AD) neuropathological measures.

**RESULTS:** Our results confirm an association of EHR-derived cognitive scores with hallmark neuropathological findings. Notably, higher neuropathological load, particularly neuritic plaques, was associated with higher cognitive burden scores in the frontal (ß=0.38, p=0.0004), parietal (ß=0.35, p=0.0008), temporal (ß=0.37, p=0. 0004) and occipital (ß=0.37, p=0.0003) lobes.

**DISCUSSION:** This proof of concept study supports the validity of NLP-based methodologies to obtain quantitative measures of RDoC clinical domains from postmortem EHR.

## 1. Introduction

Human post-mortem brain research provides a critical link between *in-vitro* studies, animal models, and human clinical studies on brain disorders. Its unique contribution is access to the human brain at the cellular and molecular levels. However, post-mortem brain research also has some limitations, including the extent of available information regarding donor demographics and, particularly, clinical phenotypes [1–5]. This latter is particularly challenging. In most cases, categorical diagnoses are derived using information gleaned from medical records, the legal-next-of-kin, and neuropathological assessments. This approach is widely used (e.g. [6,7]) and is the backbone of innumerable ground-breaking findings on the pathophysiology of a broad range of brain disorders [8]. In addition to its inherent categorical framework, other limitations, shared by the majority of studies on human studies in this field, are related to clinical phenotypic heterogeneity and overlapping comorbidities typical of many brain disorders and challenges to account for the course of disease over time. These limitations point to the need to bring human brain postmortem investigations in line with current transdiagnostic frameworks [9–14] so that the underlying pathophysiology of brain disorders can be interpreted in a more nuanced manner in the context of transdiagnostic dimensional phenotypes [15–18]. Dimensional clinical phenotypes find support in compelling evidence for substantial overlap of genetic risk as well as clinical, pathophysiological, and pharmacological features across categorical brain disorders [9–14,19,20]. Recent progress in computational sciences allows for the analysis of health-related data through machine learning, including the analysis of electronic health records (EHR) for comprehensive clinical phenotyping beyond a given diagnosis. Previous literature has focused on using machine learning methods, such as autoencoders, convolutional, and recurrent neural networks, to read clinically relevant texts and predict clinical outcomes including readmission rate, risk classifications, discharge timeline, treatment outcome as well as sub-phenotyping [21–23]. Among these deep learning approaches, natural language processing (NLP) - software designed to extract information from human-authored narrative-free text - has been widely applied to the medical field to profile various brain disorders and symptoms through algorithms ranging from speech recognition to syntax and sentiment analysis [24]. In the clinical setting, this methodology has been validated against expert annotation, formal cognitive testing, and clinical prediction tasks [25–31]. To our knowledge, it has not yet been applied to the health records from post-mortem brain donors and used in combination with neuropathological readouts.

Potential frameworks that can be applied to EHR for multidimensional clinical phenotyping include the Research Domain Criteria (RDoC) and the Hierarchical Taxonomy of Psychopathology (HiTOP) [9,10]. The National Institute of Mental Health (NIMH) developed the RDoC framework, a clinical domain-based approach with each domain designed to capture a spectrum of symptoms rooted in brain circuits and biology [11–14]. The domain-specific symptom burden can be estimated from patient medical records using NLP [28,32,33]. RDoC symptom burdens estimated from medical records by NLP have been associated with genetic variants and clinical outcomes including suicide, hospital utilization, new dementia diagnosis, and progression from dementia diagnosis to death [34–36].

We put forward that application of NLP-based methodologies to human brain postmortem studies may represent a significant step toward a more current and translatable interpretation of molecular and cellular read-outs in the context of transdiagnostic clinical domains and symptom constructs. As a first step toward assessing the feasibility and validity of this approach, we focused on Alzheimer’s Disease (AD), a disease with distinct symptoms and well-established neuropathological hallmarks. These arise from two dominant protein pathologies: Amyloid-β (Aβ), forming extracellular Aβ aggregates known as amyloid, or senile, plaques and Tau, forming intracellular Tau accumulations known as neurofibrillary tangles (NFT) and dystrophic neurites (DNs). Neuritic plaques (NP), formed by β-amyloid plaques containing DNs, are considered a pathologic hallmark of AD [37]. Clinically, AD is characterized by impaired cognition including deficits of mnestic and non-mnestic memory, judgment, and reasoning, as well as impaired visuospatial and language functions[38–40].

The aim of this study is to provide conceptual evidence for the use of NLP on post-mortem brain donor health records and the association of NLP-derived dimensional RDoC phenotypes with AD hallmark neuropathology. In line with prior evidence for an association of cognitive decline and NP burden and as a proof-of-concept study, we hypothesize that NLP-derived scores for the RDoC cognitive domain are associated with NP load.

## 2. Material and methods

### 2.1 Study Cohort

We selected 92 donors (46 males, 46 females, mean age 81.9 years, SD 9.47, range: 57 to 98 years) from the Harvard Brain and Tissue Resource Center, NIH NeuroBioBank (HBTRC/NBB) with Braak & Braak stages between 0 and 6. Post-mortem clinical and neuropathological evaluation was performed by an experienced team of a neuropathologist and two psychiatrists. Briefly, health records were reviewed by two clinicians independently and a postmortem clinical diagnosis was determined during a consensus diagnosis meeting. The neuropathological report included gross examination, macroscopic and microscopic assessment of a standard set of brain regions, as well as a semiquantitative neuritic plaque rating. Several features, including neuronal loss and presence of NFTs and neuritic plaques were assessed in the frontal (BA 3/2/1, 4, 9, 46), parietal (BA 39, 40), temporal (hippocampal formation with lateral geniculate body and tail of caudate nucleus, entorhinal cortex and anterior hippocampus), and occipital (BA 17, 18/19) cortex. Beside AD-typical neuropathological findings, donors showed age-related vascular changes and concomitant neuropathological findings that were considered inconsequential for the analysis including one donor with normal pressure hydrocephalus and hippocampal sclerosis each, seven cases with mild agyrophilic grain disease and seven cases with Lewy bodies. Clinically, donors were either neurotypical or diagnosed with AD/dementia without additional psychiatric or neurological dieases present.

In addition, the HBTRC collects extensive demographic data and clinical data included in health records and a questionnaire completed by the legal next-of-kin [41]. All medical records available for each case were digitized.

### 2.2 Ethics Statement

Tissue and medical records have been collected under the HBTRC IRB protocol 2015P002028 (McLean/MGB Institutional Review Board). Formal informed consent to donate the brain and related samples for research is obtained after death from the legal next-of-kin and documented in writing. Data about brain donors is made available to investigators in de-identified form according to HIPAA regulations.

### 2.3 Processing of Digital Records and Scoring Clinical Text for Cognition Symptom Burden

All available medical notes and records for each donors, regardless of time of creation or content, were scanned and transformed into a text file. Health records included documents covering the psychiatric or neurological status of the donor, documents pertaining to internal medicine, surgery, or other treatments, as well as administrative information. The present study used a previously described and validated NLP algorithm for quantifying estimated cognitive symptoms from narrative clinical text [28]. In brief, this method relies on recognizing a pre-specified set of symptom-related terms within the available records. The term list was developed through an iterative process of refinement seeded with lists of terms developed by a group of clinical experts, including the NIMH Research Domain Criteria Working Group. That seed was subsequently expanded through unsupervised machine learning to enhance coverage of the clinical lexicon [28]. The final cognitive symptom score is the proportion of terms that appear in any given note. The tool is implemented as freely available code for online download, including the full list of tokens as described in the initial validation publication (https://github.com/thmccoy/CQH-Dimensional-Phenotyper) [28]. Importantly, this tool was not trained to predict nor fitted against any particular outcome; rather, it was developed without regard to any particular categorical diagnosis or outcome, aiming instead to directly capture dimensions of neuropsychiatric symptomatology as dimensional rather than categorical [11]. Similarly, and as in prior work applying this NLP approach to dementia and general medical records across diagnosis, the algorithm was not trained, fitted, calibrated, modified, or otherwise biased toward the samples reported in this paper [35,42].

### 2.4 Analysis and Data Availability

The demographic characteristics of the cohort were summarized using univariate summary statistics. The primary analysis assessed the strength of association between cognitive symptoms and neuropathological findings of plaques by lobe. For each scanning method, we regressed the NLP-derived cognitive symptom burden score on the NP load of each lobe of the brain, controlling for age and sex, in a random intercept model. All analysis used R v4.2.2. Raw measurements are stored at the primary study site and can be provided upon request.

## 3. Results

Mixed effects models, adjusted for age and sex, combining NLP data with neuropathology results showed that higher NP load is associated with higher cognitive burden scores in the frontal (ß=0.38, p=0.0004), parietal (ß=0.35, p=0.0008), temporal (ß=0.37, p=0. 0004) and occipital (ß=0.37, p=0.0003) lobes (Table 1). A second confirmatory regression model comparing cognitive burden and Braak & Braak stage, controlling for age and sex, also showed a significant association between RDoC “cognition” and B&B stage of ß=0.35 (95%-CI: 0.16 – 0.54, p-value = 0.0005).

**Table 1:**
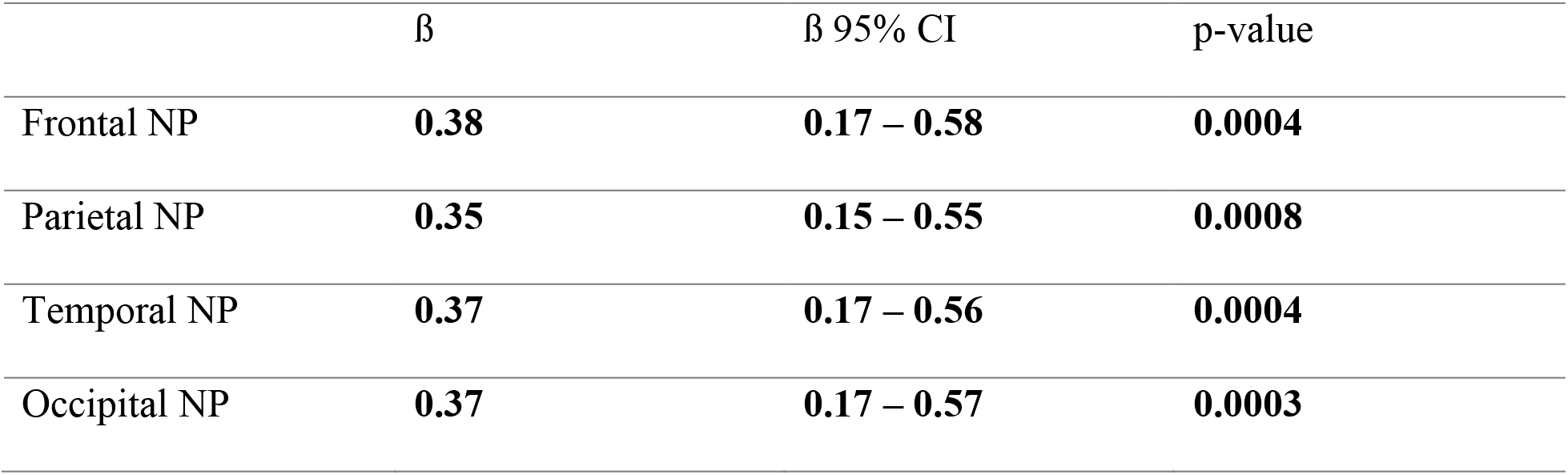
**Association between neuritic plaque load and cognitive symptom burden in mixed effect model regressing cognitive burden score on neuritic plaque load controlling for age and sex.**

## 4. Discussion

NLP methodologies, designed to obtain multidimensional phenotypic fingerprints from electronic health records, based on the NIH RDoC framework, and well-validated through expert formal cognitive testing and expert annotation [25–31]. Our goal was to validate the use of this approach in the context of human postmortem studies. Our study replicates well-established associations of cognitive decline and AD pathology, showing a strong significant correlation between pathological hallmarks of AD, i.e. NPs density and Braak and Braak stages, and cognitive deficits according to the RDoC Cognitive domain. Thus, these results offer a first proof-of-concept in support of the application of NLP algorithms based on RDoC to human postmortem studies.

A potential limitation of these studies is that the donor cohort was not followed longitudinally during the course of the disease so that we were not able to assess our results against clinical scales - health records and information from the donors’ families were obtained postmortem. Thus, clinical information on donors may vary in quality and quantity, a common challenge in postmortem studies, one that limited our options for assessment of NLP data against an accurate interpretation of clinical assessments. To address this limitation, AD-related pathological load was used in these studies as a stand-in for cognitive impairment. Although the precise relationship between aspects of AD pathology and cognitive impairment may still be under investigation, compelling evidence shows that the severity of NP and NFT load is a strong predictor of cognitive decline [43–45]. Conversely, the validity of cognitive measures derived by the NLP algorithms used for these studies is supported by extensive studies in much larger cohorts of live subjects [28,31,35]. This methodology has proven to have robust transdiagnostic predictive validity against genetic correlates as well as a broad range of clinical constructs, including agitation, risk of dementia, and risk of suicide [29,36], supporting its capacity to gain traction on difficult neuropsychiatric problems.

Transdiagnostic dimensional phenotypes are essential to investigations of the relationships between continuous symptom dimensions and cellular/molecular changes in brain disorders. In response to this challenge, efforts over the last decade have focused on overcoming the limitations inherent to categorical diagnostic approaches by establishing dimensional models based on neurobiological or behavioral phenotypes such as RDoC or HiTOP. Although these efforts are not without controversy [46], dimensional phenotyping across diagnostic entities is a critical tool needed to understand the cell-level patterns of molecular changes underlying clinical domains and symptom constructs in brain disorders. Furthermore, these approaches are needed to investigate urgent questions on the relationships between specific symptoms and underlying pathophysiological mechanisms across disorders. For instance, cognitive impairment, anxiety, and depression are shared by many brain disorders, from dementias to major depression and schizophrenia, and each is largely treated using the same pharmacological approaches. The underlying, often implicit, assumption that similar molecular, cellular, and neural circuit pathology underlies each of these symptoms across these disorders may be plausible, but as yet poorly tested. Arguably, these considerations apply to the neuropathology of dementias, as hallmark pathology, such as the impact of proteinopathies affecting tau, β-amyloid, α-synuclein, has been tested against cognition while largely neglecting co-occurring symptoms. Categorical diagnostic approaches, encompassing heterogeneous symptoms under one diagnosis, are not sufficiently nuanced to address these questions.

In conclusion, this proof-of-concept study supports the validity of NLP approaches to extract dimensional clinical phenotypic data from health records obtained postmortem from brain donors. Analyses were limited to the cognitive dimension so that hallmark AD pathology could be used as a well-established predictor of cognitive impairment. Ongoing efforts are focused on the application of these approaches to a broader range of clinical domains and brain disorders.[35].

## Acknowledgments

Brain tissue samples included in these studies were obtained from the Harvard Brain and Tissue Resource Center, an NIH NeuroBioBank site. The authors express gratitude to all brain donors and their families for their generosity. This study was funded by the National Institute of Mental Health (R01MH120991, R01MH116270). The sponsors had no role in the study design, writing of the report, or data collection, analysis, or interpretation. All authors had full access to the data and decided to submit for publication jointly.

## References

[1] McCullumsmith RE, Hammond JH, Shan D, Meador-Woodruff JH. Postmortem Brain: An Underutilized Substrate for Studying Severe Mental Illness. Neuropsychopharmacol 2014;39:65–87. https://doi.org/10.1038/npp.2013.239.

[2] Lewis DA. The Human Brain Revisited: Opportunities and Challenges in Postmortem Studies of Psychiatric Disorders. Neuropsychopharmacol 2002;26:143–54. https://doi.org/10.1016/s0893-133x(01)00393-1.

[3] Gomez-Nicola D, Boche D. Post-mortem analysis of neuroinflammatory changes in human Alzheimer’s disease. Alzheimer’s Res Ther 2015;7:42. https://doi.org/10.1186/s13195-015-0126-1.

[4] Sullivan K, Pantazopoulos H, Liebson E, Woo T-UW, Baldessarini RJ, Hedreen J, et al. Chapter 14 What can we learn about brain donors? Use of clinical information in human postmortem brain research. Handb Clin Neurology 2018;150:181–96. https://doi.org/10.1016/b978-0-444-63639-3.00014-1.

[5] Berretta S, Heckers S, Benes FM. Searching human brain for mechanisms of psychiatric disorders. Implications for studies on schizophrenia. Schizophr Res 2015;167:91–7. https://doi.org/10.1016/j.schres.2014.10.019.

[6] Levey DF, Stein MB, Wendt FR, Pathak GA, Zhou H, Aslan M, et al. Bi-ancestral depression GWAS in the Million Veteran Program and meta-analysis in >1.2 million individuals highlight new therapeutic directions. Nat Neurosci 2021;24:954–63. https://doi.org/10.1038/s41593-021-00860-2.

[7] Gelernter J, Sun N, Polimanti R, Pietrzak R, Levey DF, Bryois J, et al. Genome-wide association study of post-traumatic stress disorder reexperiencing symptoms in >165,000 US veterans. Nat Neurosci 2019;22:1394–401. https://doi.org/10.1038/s41593-019-0447-7.

[8] Wang D, Liu S, Warrell J, Won H, Shi X, Navarro FCP, et al. Comprehensive functional genomic resource and integrative model for the human brain. Science 2018;362. https://doi.org/10.1126/science.aat8464.

[9] Dalgleish T, Black M, Johnston D, Bevan A. Transdiagnostic Approaches to Mental Health Problems: Current Status and Future Directions. J Consult Clin Psych 2020;88:179–95. https://doi.org/10.1037/ccp0000482.

[10] Hengartner MP, Lehmann SN. Why Psychiatric Research Must Abandon Traditional Diagnostic Classification and Adopt a Fully Dimensional Scope: Two Solutions to a Persistent Problem. Frontiers Psychiatry 2017;8:101. https://doi.org/10.3389/fpsyt.2017.00101.

[11] Insel T, Cuthbert B, Garvey M, Heinssen R, Pine DS, Quinn K, et al. Research Domain Criteria (RDoC): Toward a New Classification Framework for Research on Mental Disorders. Am J Psychiat 2010;167:748–51. https://doi.org/10.1176/appi.ajp.2010.09091379.

[12] Cuthbert BN. The RDoC framework: facilitating transition from ICD/DSM to dimensional approaches that integrate neuroscience and psychopathology. World Psychiatry 2014;13:28–35. https://doi.org/10.1002/wps.20087.

[13] Insel TR, Cuthbert BN. Medicine. Brain disorders? Precisely. Science 2015;348:499–500. https://doi.org/10.1126/science.aab2358.

[14] Carcone D, Ruocco AC. Six Years of Research on the National Institute of Mental Health’s Research Domain Criteria (RDoC) Initiative: A Systematic Review. Front Cell Neurosci 2017;11:46. https://doi.org/10.3389/fncel.2017.00046.

[15] Reininghaus U, Böhnke JR, Chavez-Baldini U, Gibbons R, Ivleva E, Clementz BA, et al. Transdiagnostic dimensions of psychosis in the Bipolar-Schizophrenia Network on Intermediate Phenotypes (B-SNIP). World Psychiatry 2019;18:67–76. https://doi.org/10.1002/wps.20607.

[16] Parkes L, Tiego J, Aquino K, Braganza L, Chamberlain SR, Fontenelle LF, et al. Transdiagnostic variations in impulsivity and compulsivity in obsessive-compulsive disorder and gambling disorder correlate with effective connectivity in cortical-striatal-thalamic-cortical circuits. Neuroimage 2019;202:116070. https://doi.org/10.1016/j.neuroimage.2019.116070.

[17] Grisanzio KA, Goldstein-Piekarski AN, Wang MY, Ahmed APR, Samara Z, Williams LM. Transdiagnostic Symptom Clusters and Associations With Brain, Behavior, and Daily Function in Mood, Anxiety, and Trauma Disorders. Jama Psychiat 2017;75:201. https://doi.org/10.1001/jamapsychiatry.2017.3951.

[18] Cowan HR, Mittal VA. Transdiagnostic Dimensions of Psychiatric Comorbidity in Individuals at Clinical High Risk for Psychosis: A Preliminary Study Informed by HiTOP. Frontiers Psychiatry 2021;11:614710. https://doi.org/10.3389/fpsyt.2020.614710.

[19] Romero C, Werme J, Jansen PR, Gelernter J, Stein MB, Levey D, et al. Exploring the genetic overlap between twelve psychiatric disorders. Nat Genet 2022;54:1795–802. https://doi.org/10.1038/s41588-022-01245-2.

[20] Hindley G, Frei O, Shadrin AA, Cheng W, O’Connell KS, Icick R, et al. Charting the Landscape of Genetic Overlap Between Mental Disorders and Related Traits Beyond Genetic Correlation. Am J Psychiat 2022;179:833–43. https://doi.org/10.1176/appi.ajp.21101051.

[21] Rajkomar A, Oren E, Chen K, Dai AM, Hajaj N, Hardt M, et al. Scalable and accurate deep learning with electronic health records. Npj Digital Medicine 2018;1:18. https://doi.org/10.1038/s41746-018-0029-1.

[22] Solares JRA, Raimondi FED, Zhu Y, Rahimian F, Canoy D, Tran J, et al. Deep learning for electronic health records: A comparative review of multiple deep neural architectures. J Biomed Inform 2020;101:103337. https://doi.org/10.1016/j.jbi.2019.103337.

[23] Lage I, Jr THM, Perlis RH, Doshi-Velez F. Efficiently identifying individuals at high risk for treatment resistance in major depressive disorder using electronic health records. J Affect Disorders 2022;306:254–9. https://doi.org/10.1016/j.jad.2022.02.046.

[24] Hecker P, Steckhan N, Eyben F, Schuller BW, Arnrich B. Voice Analysis for Neurological Disorder Recognition–A Systematic Review and Perspective on Emerging Trends. Frontiers Digital Heal 2022;4:842301. https://doi.org/10.3389/fdgth.2022.842301.

[25] Perlis RH, Iosifescu DV, Castro VM, Murphy SN, Gainer VS, Minnier J, et al. Using electronic medical records to enable large-scale studies in psychiatry: treatment resistant depression as a model. Psychol Med 2012;42:41–50. https://doi.org/10.1017/s0033291711000997.

[26] McCoy TH, Castro VM, Rosenfield HR, Cagan A, Kohane IS, Perlis RH. A clinical perspective on the relevance of research domain criteria in electronic health records. Am J Psychiat 2015;172:316–20. https://doi.org/10.1176/appi.ajp.2014.14091177.

[27] Perlis RH, Fava M, McCoy TH. Can electronic health records revive central nervous system clinical trials? Mol Psychiatr 2019;24:1096–8. https://doi.org/10.1038/s41380-018-0278-z.

[28] McCoy TH, Yu S, Hart KL, Castro VM, Brown HE, Rosenquist JN, et al. High Throughput Phenotyping for Dimensional Psychopathology in Electronic Health Records. Biol Psychiat 2018;83:997–1004. https://doi.org/10.1016/j.biopsych.2018.01.011.

[29] McCoy TH, Castro VM, Roberson AM, Snapper LA, Perlis RH. Improving Prediction of Suicide and Accidental Death After Discharge From General Hospitals With Natural Language Processing. Jama Psychiat 2016;73:1064. https://doi.org/10.1001/jamapsychiatry.2016.2172.

[30] Barroilhet SA, Pellegrini AM, McCoy TH, Perlis RH. Characterizing DSM-5 and ICD-11 personality disorder features in psychiatric inpatients at scale using electronic health records. Psychol Med 2020;50:2221–9. https://doi.org/10.1017/s0033291719002320.

[31] Hart KL, Pellegrini AM, Forester BP, Berretta S, Murphy SN, Perlis RH, et al. Distribution of agitation and related symptoms among hospitalized patients using a scalable natural language processing method. Gen Hosp Psychiat 2021;68:46–51. https://doi.org/10.1016/j.genhosppsych.2020.11.003.

[32] McCoy TH, Castro VM, Rosenfield HR, Cagan A, Kohane IS, Perlis RH. A Clinical Perspective on the Relevance of Research Domain Criteria in Electronic Health Records. Am J Psychiat 2015;172:316–20. https://doi.org/10.1176/appi.ajp.2014.14091177.

[33] Filannino M, Stubbs A, Uzuner Ö. Corrigendum to “Symptom severity prediction from neuropsychiatric clinical records: Overview of 2016 CEGS N-GRID shared tasks Track 2” [J Biomed Inform. 2017 Nov;75S:S62-S70]. J Biomed Inform 2018;85:204. https://doi.org/10.1016/j.jbi.2018.08.015.

[34] McCoy TH, Pellegrini AM, Perlis RH. Differences among Research Domain Criteria score trajectories by Diagnostic and Statistical Manual categorical diagnosis during inpatient hospitalization. Plos One 2020;15:e0237698. https://doi.org/10.1371/journal.pone.0237698.

[35] McCoy TH, Han L, Pellegrini AM, Tanzi RE, Berretta S, Perlis RH. Stratifying risk for dementia onset using large-scale electronic health record data: a retrospective cohort study. Alzheimer’s Dementia 2019;16:531–40. https://doi.org/10.1016/j.jalz.2019.09.084.

[36] McCoy TH, Pellegrini AM, Perlis RH. Research Domain Criteria scores estimated through natural language processing are associated with risk for suicide and accidental death. Depress Anxiety 2019;36:392–9. https://doi.org/10.1002/da.22882.

[37] Braak H, Braak E. Demonstration of amyloid deposits and neurofibrillary changes in whole brain sections. Brain Pathol 1991;1:213–6. https://doi.org/10.1111/j.1750-3639.1991.tb00661.x.

[38] McKhann GM, Knopman DS, Chertkow H, Hyman BT, Jack CR, Kawas CH, et al. The diagnosis of dementia due to Alzheimer’s disease: Recommendations from the National Institute on Aging-Alzheimer’s Association workgroups on diagnostic guidelines for Alzheimer’s disease. Alzheimer’s Dementia 2011;7:263–9. https://doi.org/10.1016/j.jalz.2011.03.005.

[39] Dubois B, Hampel H, Feldman HH, Scheltens P, Aisen P, Andrieu S, et al. Preclinical Alzheimer’s disease: Definition, natural history, and diagnostic criteria. vol. 12. 2016. https://doi.org/10.1016/j.jalz.2016.02.002.

[40] Jack CR, Bennett DA, Blennow K, Carrillo MC, Dunn B, Haeberlein SB, et al. NIA-AA Research Framework: Toward a biological definition of Alzheimer’s disease. Alzheimer’s Dementia 2018;14:535–62. https://doi.org/10.1016/j.jalz.2018.02.018.

[41] Sullivan K, Pantazopoulos H, Liebson E, Woo TUW, Baldessarini RJ, Hedreen J, et al. What can we learn about brain donors? Use of clinical information in human postmortem brain research. Handb Clin Neurology 2018;150:181–96. https://doi.org/10.1016/b978-0-444-63639-3.00014-1.

[42] Hart KL, Perlis RH, McCoy TH. Mapping of Transdiagnostic Neuropsychiatric Phenotypes Across Patients in Two General Hospitals. J Acad Consult Psychiatry 2021;62:430–9. https://doi.org/10.1016/j.jaclp.2021.01.002.

[43] Nelson PT, Alafuzoff I, Bigio EH, Bouras C, Braak H, Cairns NJ, et al. Correlation of Alzheimer Disease Neuropathologic Changes With Cognitive Status: A Review of the Literature. J Neuropathology Exp Neurology 2012;71:362–81. https://doi.org/10.1097/nen.0b013e31825018f7.

[44] Nelson PT, Jicha GA, Schmitt FA, Liu H, Davis DG, Mendiondo MS, et al. Clinicopathologic correlations in a large Alzheimer disease center autopsy cohort: Neuritic plaques and neurofibrillary tangles “do count” when staging disease severity. J Neuropathology Exp Neurology 2007;66:1136–46. https://doi.org/10.1097/nen.0b013e31815c5efb.

[45] Hyman BT, Phelps CH, Beach TG, Bigio EH, Cairns NJ, Carrillo MC, et al. National Institute on Aging–Alzheimer’s Association guidelines for the neuropathologic assessment of Alzheimer’s disease. Alzheimer’s Dementia 2012;8:1–13. https://doi.org/10.1016/j.jalz.2011.10.007.

[46] Ross CA, Margolis RL. Research Domain Criteria: Strengths, Weaknesses, and Potential Alternatives for Future Psychiatric Research. Mol Neuropsychiatry 2019;5:218–36. https://doi.org/10.1159/000501797.

